# DNA origami nanostructures as a tool for the targeted destruction of bacteria

**DOI:** 10.1101/837252

**Authors:** Ioanna Mela, Pedro P. Vallejo-Ramirez, Stanislaw Makarchuk, Graham Christie, David Bailey, Robert. M. Henderson, Hiroshi Sugiyama, Masayuki Endo, Clemens F. Kaminski

## Abstract

Antibiotic resistance is a growing worldwide human health issue that is now rendering us vulnerable once again to infections that have been treatable for decades. Various approaches have been proposed in an effort to overcome this threat and effectively treat bacterial infections. We use a DNA origami nanostructure, functionalized with aptamers, as a vehicle for delivering the antibacterial peptide lysozyme in a specific and efficient manner, in order to destroy bacterial targets. We test the system against Gram-positive (*Bacillus subtilis*) and Gram - negative (*Escherichia coli*) targets. We use direct stochastic optical reconstruction microscopy (dSTORM) and atomic force microscopy (AFM) to characterize the DNA origami nanostructures and structured illumination microscopy (SIM) to assess the binding of origami to the bacteria. We show that treatment with lysozyme-functionalized origami slows bacterial growth more effectively than treatment with free lysozyme. Our study introduces DNA origami as a tool in the fight against antibiotic resistance, and our results demonstrate the specificity and efficiency of the nanostructure as a drug delivery vehicle.

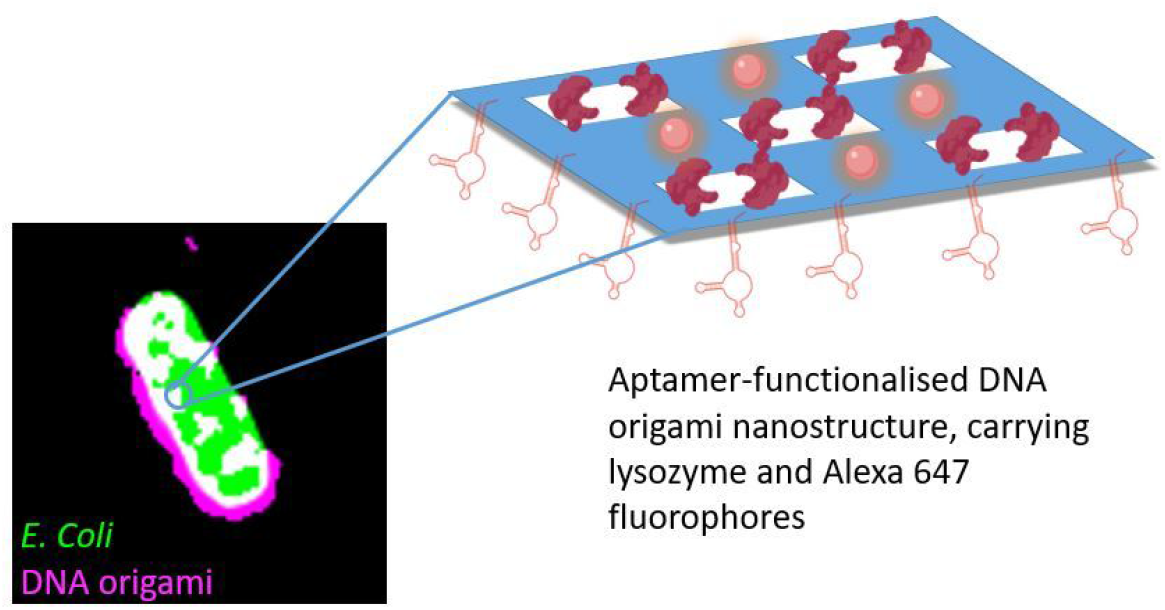

---

The rate at which bacteria (especially Gram-negative pathogens) are developing resistance to antimicrobial agents is higher than the rate at which new antibiotics are being developed, increasing the risk that untreatable infections will become widespread. Resistance of *Escherichia coli* and *Klebsiella pneumoniae* to third-generation cephalosporins substantially increased in the EU between 2012 and 2015 (to >50% resistance of *K. pneumoniae* in many countries) and resistance of *Pseudomonas aeruginosa* and *Acinetobacter* spp. to multiple antimicrobial groups has also become common^1^. According to the World Health Organization, failure of last-resort treatments for gonorrhea has been confirmed in at least ten countries, and failure of treatment for *E. coli* urinary tract infections, *S. aureus* infections and *Enterobacteriaceae* infections has been reported widely around the world^2^. Therefore, alternative antimicrobial strategies are needed urgently.

Several novel materials, including metal-organic frameworks^3^, antimicrobial peptides^4–6^, nanoparticles^7–9^ and combinations of these^10–12^, have shown promise for new antimicrobial strategies, but problems persist. For example, metal-based materials have low stability and/or can be highly toxic to mammalian cells^13–16^. Similarly, antimicrobial peptides are promising, but methods for their targeted delivery are still lacking. In this study, we explore the potential of DNA origami as a vehicle for delivering active antimicrobial components in a target-specific and efficient manner.

DNA origami structures are two- or three-dimensional nanostructures made by exploiting the base-pairing property of DNA^17^. A large number (150-200) of oligonucleotides — referred to as staples — are used to fold the DNA into a pre-designed conformation and hold it together, and these staples can be functionalized to carry various payloads^18,19^. Previous studies have shown that DNA origami has excellent biocompatibility, triggers no immune response and stays intact in vivo for at least 48 h^20,21^, making it an ideal candidate material for the manufacturing of highly specific drug delivery vehicles. For example, DNA origami has been used to target active therapeutic molecules to eukaryotic cancer cells, resulting in the death of these cells^21,22^. However, there are no reports on the use of DNA origami to target bacterial cells.

In this study, we used DNA origami as a vehicle for targeted delivery of the antimicrobial enzyme lysozyme to two different bacteria *in vitro*. With only minor modifications, the same nanostructure was functionalized to target Gram-positive and Gram-negative bacteria with high specificity. We synthesized and used a previously reported DNA origami nanostructure^23^, which consists of a frame containing five ‘wells’ to carry molecular payloads, and functionalized it with aptamers designed to target *E. coli* and *B. subtilis* bacterial strains^24^ (Figure 1). Successful formation of the DNA frames was verified with atomic force microscopy (AFM). Approximately 80% of the frames had formed as designed. (Figure 2a). The mean measured length and width of the frames (~100 × 100 nm; n = 105) agreed with those of the frame design. The wells (Figure 2.a, b) measured ~20 × 15 nm.

**Figure 1.**
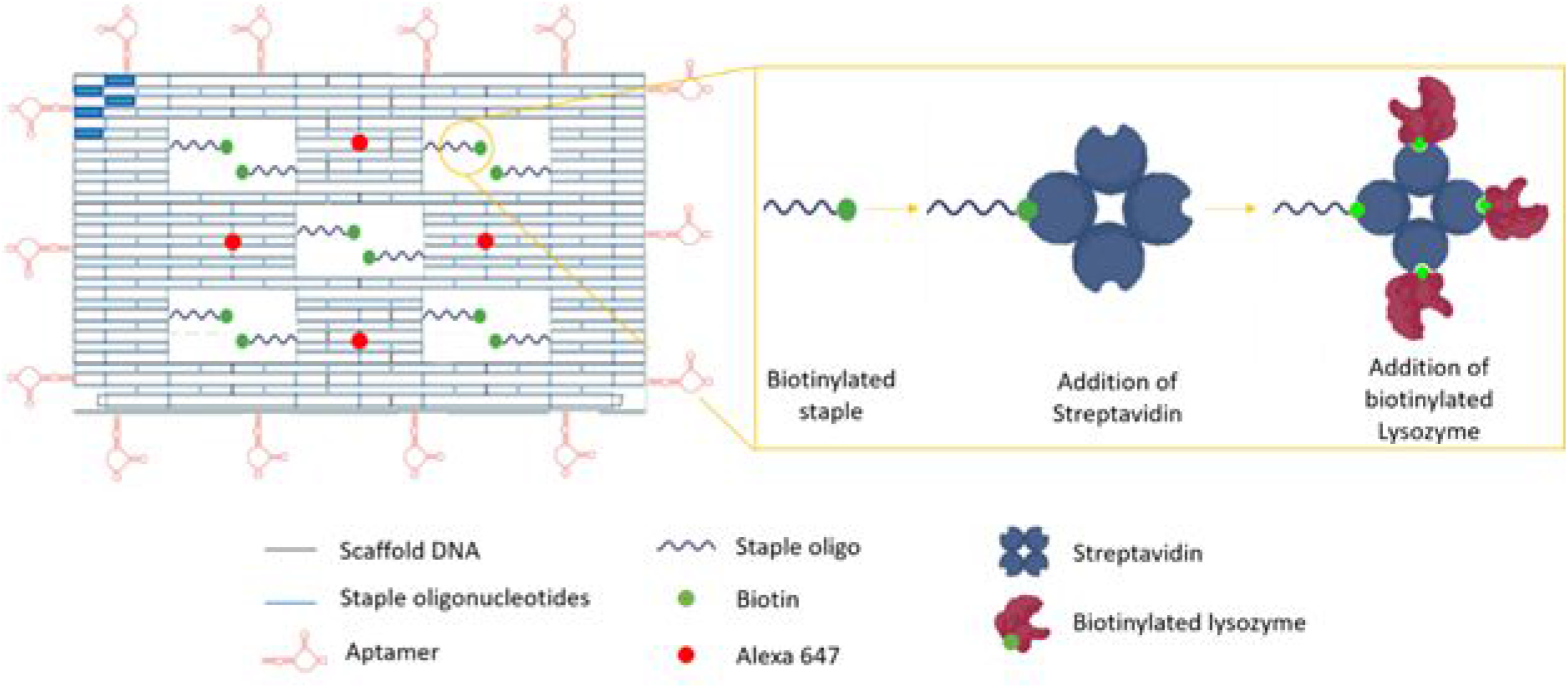
Schematic representation of the DNA origami nanostructure. (left). Each of the “wells” carries two biotinylated staples for the attachment of streptavidin and, subsequently, biotinylated lysozyme (right). Fourteen aptamers (in red), hybridized with staples at the four sides of the DNA origami drive the attachment of the nanostructures to the bacterial targets. Four Alexa 647 molecules (red circles) act as detection beacons for the nanostructure. The five blue bars at the top left corner of the nanostructure represent hairpins, used for orientation.

**Figure 2.**
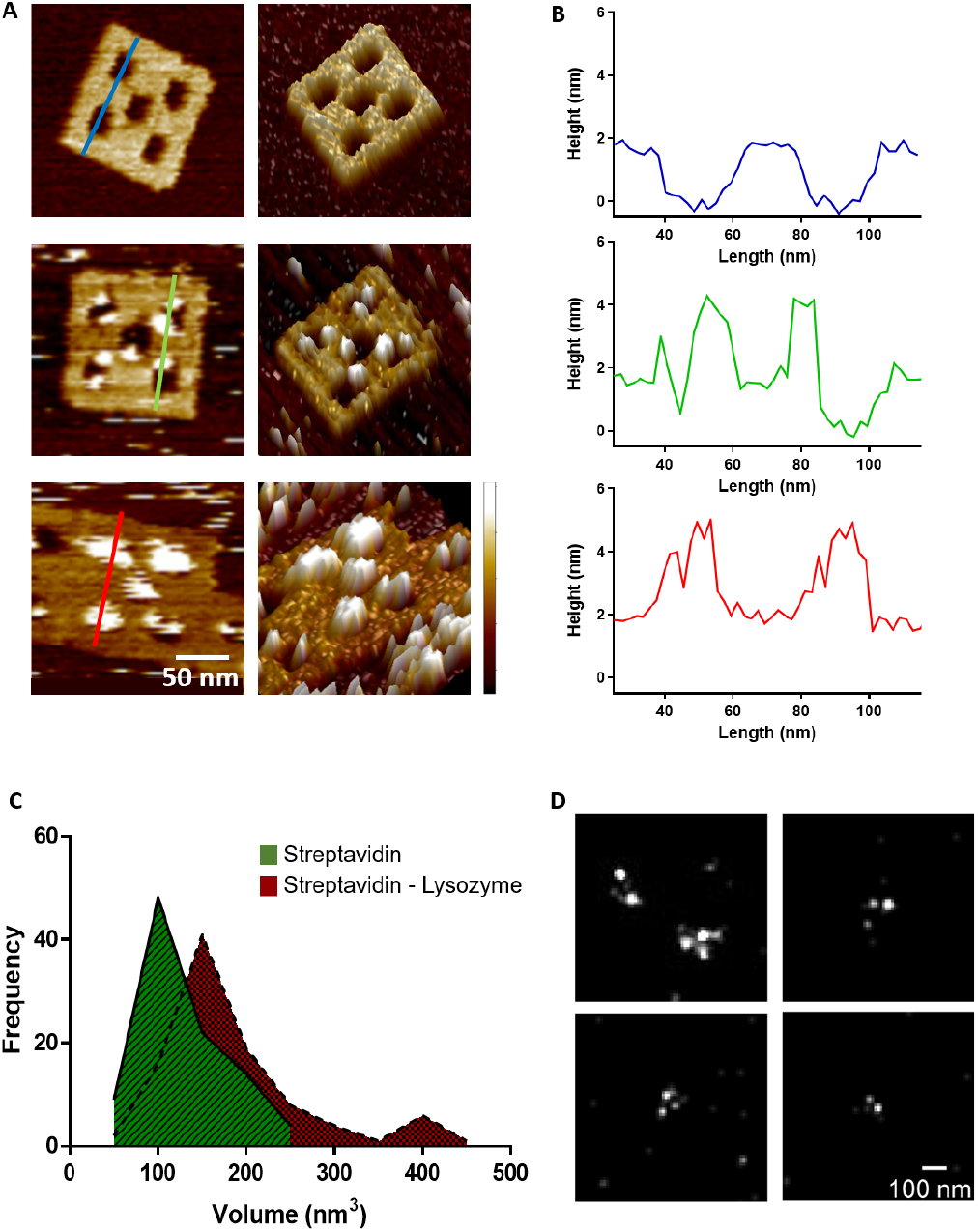
Characterization of the DNA nanostructure. **2.a** Two-dimensional (left) and three-dimensional atomic force microscopy images of DNA origami nanostructures before any incubation (top), after incubation with streptavidin (middle) and after successive incubations with streptavidin and biotinylated lysozyme (bottom) (Height scale 0-3.5 nm, from darker to lighter).**2.b** Cross-sections of the nanostructures shown in a show the change height at the wells before any incubation (blue line), after incubation with streptavidin (green line) and after successive incubations with streptavidin and biotinylated lysozyme (red line).**2.c** Volume measurements of the particles bound to the DNA origami structures after incubation with streptavidin alone (green) and streptavidin and biotinylated lysozyme (red). The volume shift corresponds to two lysozyme molecules. **2.d** dSTORM super-resolution microscopy images of Alexa 647 fluorophores in the DNA origami nanostructures, demonstrating their successful incorporation.

The design of the frames incorporated three different functionalizations. The first was the inclusion of ten biotinylated staples (two in each well – detailed sequences in SI) to enable attachment of biotin-tagged lysozyme to the frames. To attach the biotinylated lysozyme, we exploited the strong and efficient binding between biotin and streptavidin and the tetrameric structure of streptavidin that enables it to bind to four biotin molecules simultaneously. The biotin was attached to the oligonucleotide staples via a five-base linker to provide flexibility for the attachment of the streptavidin/lysozyme complex. To confirm successful attachment of lysozyme to the DNA origami frames, we used AFM to measure the volumes of molecules that were bound to the frames after incubation with streptavidin alone and with streptavidin and biotinylated lysozyme (Figure 2.c). The mean volume of bound molecules after incubation with streptavidin alone was ~110 nm^3^, which corresponds to the theoretical volume of streptavidin. The mean volume of bound molecules after incubation with streptavidin and biotinylated lysozyme was ~160 nm^3^, corresponding to the expected volume when 1–2 lysozyme molecules are bound to each streptavidin tetramer. Three or four wells per frame were occupied by streptavidin/lysozyme complexes.

The second functionalization was the inclusion of four fluorophore (Alexa 647) molecules to enable detection of the nanostructures with fluorescence microscopy. We confirmed successful incorporation of the Alexa 647 functionalized staples with dSTORM super-resolution microscopy. A mean of three fluorophores were observed to be incorporated into each structure (n=50) (Figure 2.d).

The measured distance between the single fluorophores was 62.3 +/− 17 nm (n=40), agreeing with the theoretical distance between the Alexa 647 molecules on the nanostructure.

The third functionalization was the incorporation of aptamers around the edges of the frame. Previous studies have shown that aptamers effectively and selectively bind to bacterial targets^24,25^. To ensure effective aptamer-driven binding of the DNA origami to bacterial targets, we incorporated 14 aptamers in each nanostructure (Figure 1). We used aptamers that are 40 bases long and can bind to both Gram-positive and Gram-negative bacteria^24^. Successful binding of the DNAorigami nanostructures to Gram-negative (*E. coli*) and Gram-positive (*B. subtilis*) bacterial strains was confirmed with Structured Illumination Microscopy (SIM). We used *E. coli* BL21(DE3) that expresses GFP and *B. subtilis* that was stained with Nile Red dye. Expression of GFP is visible within the whole bacterial cell, while Nile red, a lipophilic dye, stains only the outer membrane of the bacterium. As a negative control, we used *Lactococcus lactis* NZ9000 cells, which were also stained with Nile Red.

Bacteria were incubated with DNA origami, and SIM was used to visualize the bacteria and the Alexa 647-labelled DNA origami in each sample (Figure 3.a,b). The surface area of bacteria covered by DNA origami was measured by evaluating the fluorescence overlap. We observed that 83% ± 5% of the E. coli bacteria (n=825) had some degree of DNA origami decoration, while 72% ± 19% of the B. subtilis population (n= 750) was decorated with DNA origami. Interestingly, in both strains, the average area of the bacterium covered by DNA origami, was ~ 20% (n= 825 for E. coli and n= 750 for B. subtilis, Figure 3.c). The observed coverage was achieved with a DNA origami concentration of ~10 nM, indicating that the nanostructures have high affinity for the bacterial targets. Binding of DNA origami with no aptamers was minimal (less than 2% area covered); the binding observed, probably resulted from electrostatic interactions (Figure 3.d,e). Similarly, negligible binding was observed with use of L. lactis, to which aptamers cannot bind (Figure 3.f).

**Figure 3.**
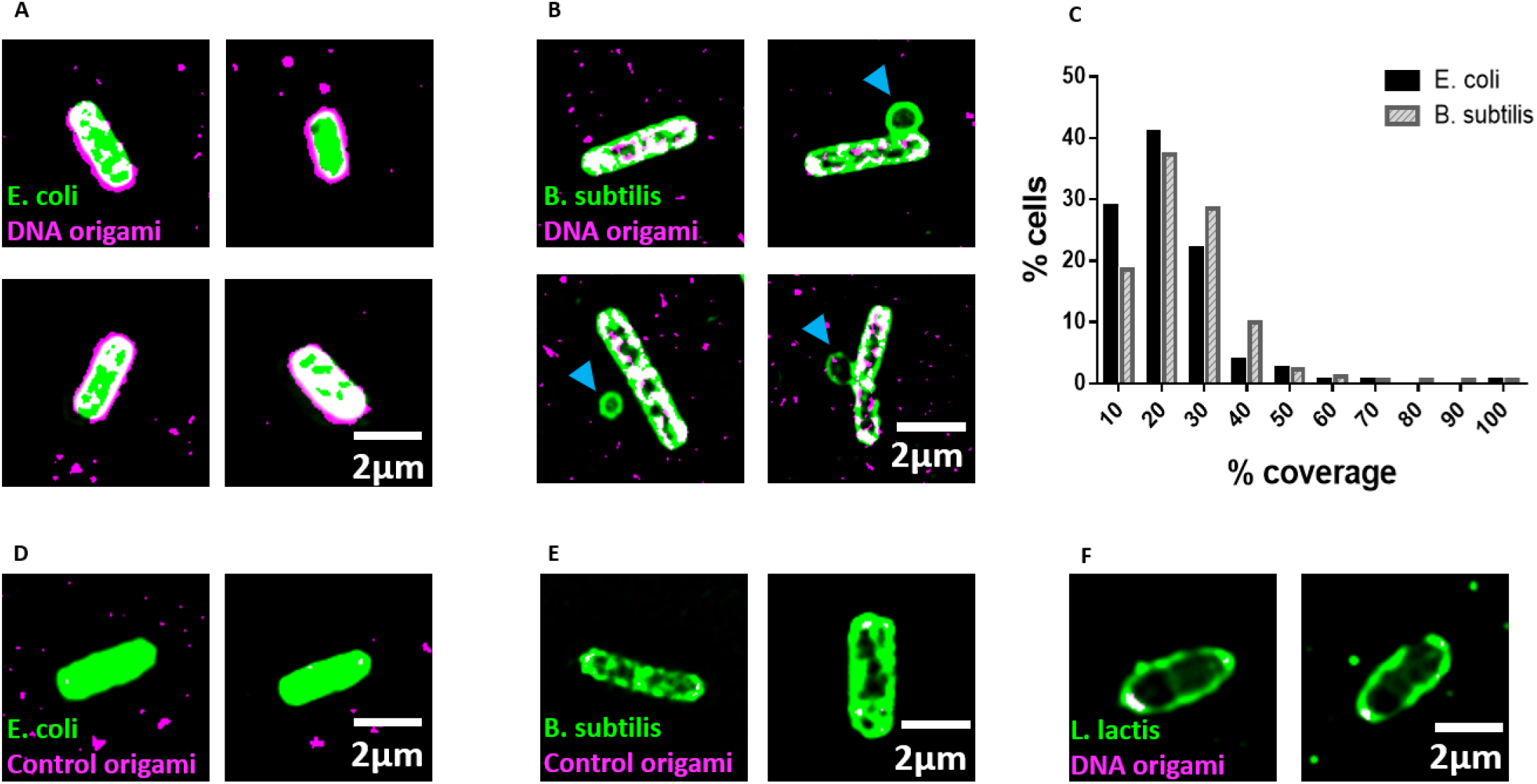
Structured Illumination microscopy of E. coli and B. subtilis. **3.a** Structured illumination microscopy (SIM) image demonstrating that DNA origami binds to *E. coli*. DNA origami is magenta and GFP-expressing *E. coli* are green, while the overlapping area are in white. **3.b** SIM imaging demonstrating that DNA origami binds to B. subtilis. DNA origami is magenta and *B. subtilis* green, with overlapping areas in white. **3.c** The mean coverage for *E. coli* is *18.6* %, and 22.5% for *B. subtilis*. **3d, e** *E. coli* (d) and *B. subtilis* (e) that were incubated with DNA origami that did not carry aptamers. Minimal binding is visible, showing that aptamers are necessary for successful binding. **3.f** SIM image of *L. lactis* incubated with DNA origami. The nanostructures do not bind this bacterial strain.

We used growth assays for *E. coli* and *B. subtilis* to investigate how free lysozyme, plain DNA origami and DNA origami carrying lysozyme affected bacterial growth over 16 hours. To extract the growth rate in each condition, we fitted the growth curves with a modified Gompertz growth equation^26^

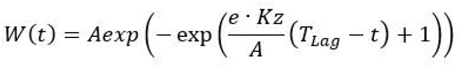

Where k_z_ is the absolute growth rate (i.e. tangent to the curve) and T_Lag_ represents the time between recovery of the microbial population from being transferred to a new habitat and the occurrence of substantial cell division. In the present case, the dependent variable W(t) represents the change in OD_600_ as a function of time. The advantage of this re-parameterisation is that the growth rate coefficient (Kz) constitutes the absolute growth rate at inflection, and that A (the upper asymptote) does not affect this parameter^27^. Nine individual growth curves were analyzed in each condition.

The growth of Gram-negative bacteria (*E. coli*) (Figure 4.a,b) was not significantly affected by free lysozyme (300 nM), as expected^28,29^. However, their growth was significantly slowed by the presence of aptamer-functionalized DNA origami loaded with lysozyme. Previous work has shown that targeted, localized delivery of the enzyme to the bacteria increases its efficiency against *E.coli*^30^ and we observe the same effect in the present study. Interestingly, we also observed significant reduction in the growth rate of *E. coli* in the presence of plain DNA origami (aptamer-functionalized but without any active payload). This reduction in the growth rate could indicate that the binding of the nanostructures to those bacteria interferes with their ability to divide and grow efficiently. Moreover, *E. coli* grown in the presence of DNA origami as well as those grown in the presence of DNA origami carrying lysozyme, also show a lower upper asymptote than the control sample. *E. coli* grown in the presence of DNA origami that does not carry aptamers were not affected (Figure S.3), indicating further that it is the binding of the nanostructures onto the bacterial targets that slows the growth rate. Identifying the exact binding sites of aptamer-functionalised DNA origami on the bacterial surface and how targeting different sites affects bacterial growth are interesting questions for future work.

**Figure 4.**
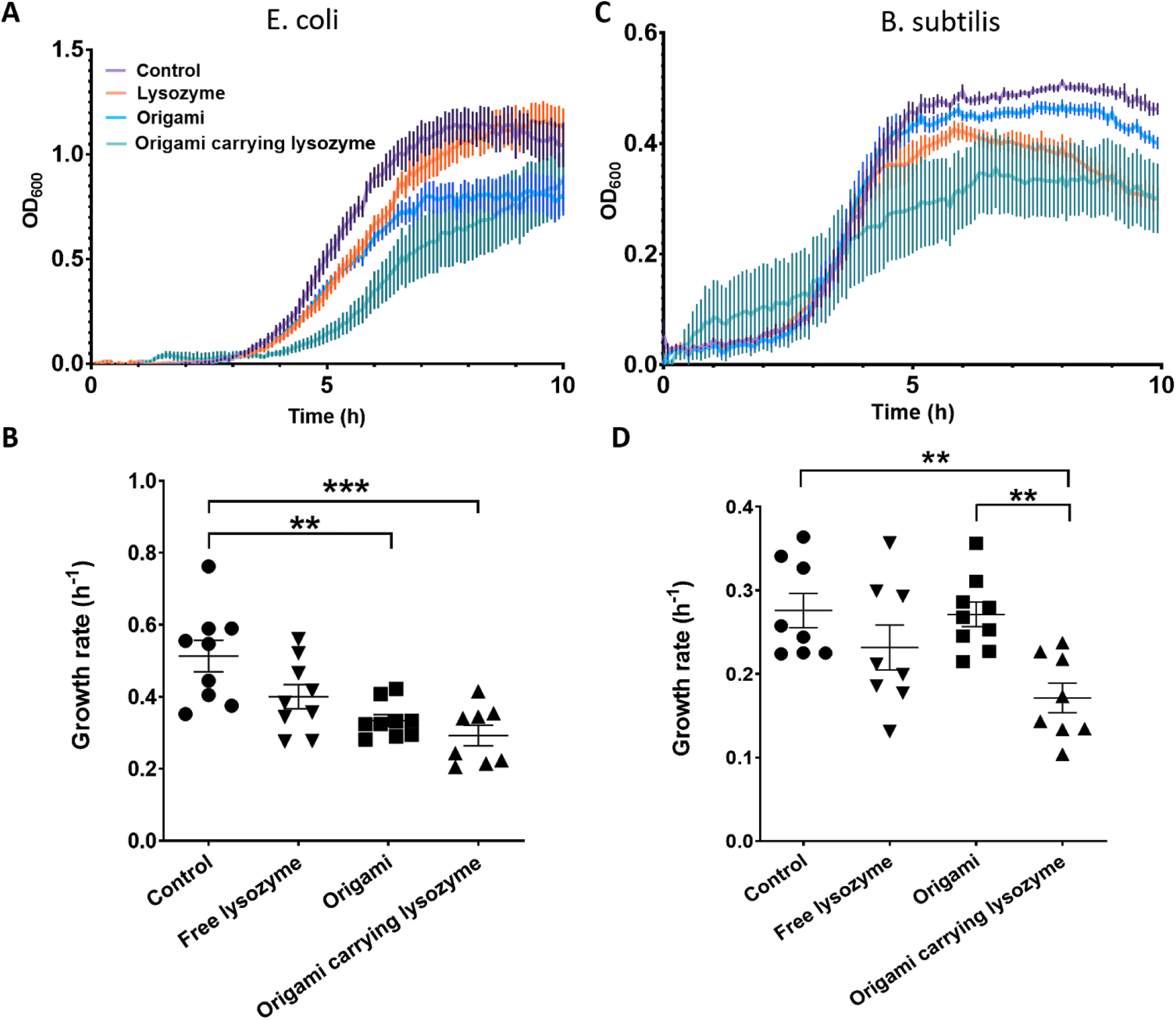
Bacterial growth analysis. **4.a** Averaged growth curves for *E.coli* (n=9) show that both the growth rate and the maximum population density are affected by the presence of DNA origami and of DNA origami carrying lysozyme in the culture medium. **4.b** Averaged growth curves for *B.subtilis* (n=9) show that both the growth rate and the maximum population density are affected by the presence of DNA origami carrying lysozyme in the culture medium. **4.c** Growth rate analysis for *E.coli* shows that the growth rate is reduced in the presence of DNA origami that carries lysozyme and in the presence of plain DNA origami but not in the presence of free lysozyme. **4.d** Growth rate analysis for *B. subtilis* shows that DNA origami carrying lysozyme significantly reduces the growth rate of the bacteria. All the origami nanostructures used in those experiments were functionalized with aptamers. Error bars in each graph represent the standard error of the mean

The growth of the Gram positive bacteria (*B. subtilis*) (Figure 4.c,d) was also significantly slowed by the presence of aptamer-functionalised DNA origami carrying lysozyme (300nM), while it was reduced, but not significantly by the presence of free lysozyme. However, plain, aptamer-functionalised DNA origami did not affect the growth of *B. subtilis* in the way it affected growth of *E. coli*. This observation leads us to believe that the precise nature of the interaction between the aptamer-derivatized nanostructures and the bacterial surface directly influences the effects of the nanostructures themselves on bacterial growth. The exact nature of this interaction is therefore an important area for future investigation and could be exploited to develop highly selective and potent antibacterial DNA nanostructures. In the *B. subtilis* sample, we noticed the presence of minicells, (blue arrows in figure 3.b). Both Gram-positive and Gram-negative bacteria have the ability to form minicells but little is known about why this happens. A recent study^31^ suggests that minicells could act as a “damage disposal” mechanism for proteins damaged by antibiotics.

To assess the effect of functionalized DNA origami on mammalian cells, we incubated COS-7 cells with DNA origami carrying lysozyme, as well as with free lysozyme. No significant effects were observed in the viability of the cells, indicating that DNA origami is a promising candidate for drug delivery *in vivo* (Figure S.4).

To conclude, we have developed a platform for bacterial targeting based on the combination of DNA origami and aptamer nanotechnology. Our DNA nanostructures can bind to designated bacterial targets and deliver the antibacterial enzyme lysozyme to slow bacterial growth. Targeted and localized delivery of multiple lysozyme molecules per bacterial cell reduces the quantity of active agent required to achieve a given antibacterial effect. Our study opens the way for the use of DNA origami as a tool in the fight against antibiotic resistance, allowing for precise pathogen targeting and for the delivery of individual or combined antimicrobial compounds. The system can be easily adapted to carry appropriate payloads for various targets, making it an attractive option for antimicrobial drug delivery. Moreover, our aptamer-derivatised origami has the potential to target and block specific targets on the bacterial surface, thus inhibiting crucial bacterial functions. In that way, a “double-hit” approach can be achieved, where the bacterium is already at a disadvantage, making it easier to destroy with antimicrobial agents

## ASSOCIATED CONTENT

DNA origami nanostructures as a tool for the targeted destruction of bacteria

**Supporting Information**. (PDF)

## Author Contributions

IM designed and performed the experiments, analysed the data and wrote the manuscript. PVR performed the dSTORM imaging and data analysis. SM wrote MATLAB scripts and helped with the data analysis. GC provided bacterial strains and contributed to data interpretation. RMH provided resources and HS and ME provided resources and contributed to the experimental design. CFK revised the manuscript. All authors have given approval to the final version of the manuscript.

## ACKNOWLEDGMENTS

We thank Dr Ana Fernandez Villegas for help with the mammalian cell culture. We also thank Professor Ulrich Keyser for his interest in the project and his help to critically assess experimental procedures and Professor Takashii Morii for his insightful input. We thank Dr Colin Hockings, Dr Katharina Scherer and Maria Zacharopoulou for valuable discussions on the project. C.F.K. acknowledges funding from the Engineering and Physical Sciences Research Council, EPSRC (EP/ H018301/1, EP/L015889/1); Wellcome Trust (089703/Z/ 09/Z, 3-3249/Z/16/Z); Medical Research Council MRC (MR/K015850/1, MR/K02292X/1); MedImmune; and Infinitus (China), Ltd. and H.S. acknowledges funding from JSPS KAKENHI (Grant Number JP16H06356).

## ABBREVIATIONS

AFM: atomic force microscopy
SIM: structured illumination microscopy
dSTORM: direct stochastic optical reconstruction microscopy

