## Supplementary information for "DNA origami nanostructures as a tool for the targeted destruction of bacteria"

#### Table of contents

**Section 1. Synthesis and characterisation of DNA origami nanostructures**

**Section 2. Imaging of the nanostructures**

**Section 3. Bacterial strains imaging**

**Section 4. Bacterial growth curves**

**Section 5. Cell viability assay**

### Section 1. Synthesis of DNA origami nanostructures

#### 1.1 Synthesis of DNA origami nanostructures

We used M13mp18 single-stranded DNA as scaffold (New England Biolabs) and complementary oligonucleotides (Integrated DNA Technologies) to synthesise DNA origami nanostructures. 159 complementary oligos, as described by Yoshidome *et al*[1]. fully occupy the whole length of the scaffold. The sequences of the oligonucleotides are listed in Table S-1, in a 5' to 3' orientation.

| Sequence Name | Sequence | Bases |
|---|---|---|
| 5wf-001 1A-hp | TAA AAA TAC CGA ACG ACC TAA AAC TCC TCT TTT GAG GAA CAA GTT TTC TTG TAT CGC CAT TTT GCA GAT TC | 71 |
| 5wf-002 1B-hp | ACC AGT CAT GGA TTA TTC CTC TTT TGA GGA ACA AGT TTT CTT GTT TAC ATT GTT TTA TTA GTA A | 64 |
| 5wf-003 1C-hp | TAA CAT CAT AGC AAT ATC CTC TTT TGA GGA ACA AGT TTT CTT GTC TTC TTT GTT TTG CCA GAA T | 64 |
| 5wf-004 1DE | CCT GAG AAT AGA CAG GAA CGG TAC TTT TTG CTT TGA CGA GCA CGG GGC GCG TAC TAT GGT TTT TGC GGG CGC | 72 |
| 5wf-005 1FG | TAG GGC GCG AAG AAA GCG AAA GGA TTT TAT CGG CAA AAT CCC TTT GAT GGT GGT TCC GAA TTT TCC GCT TTC | 72 |
| 5wf-006 1HI | CAG TCG GGG CGT TGC GCT CAC TGC TTT TCG TTG TAA AAC GAC GGG TTT TCC CAG TCA CGA TTT TAG GGG ACG | 72 |
| 5wf-007 1JK | ACG ACA GTT GCA TCT GCC AGT TTG TTT TAG CCC CAA AAA CAG GAG GTT GAT AAT CAG AAA TTT TTC ATT GCC | 72 |
| 5wf-008 1LM | TGA GAG TCT ACA AAG GCT ATC AGG TTT TGT CAA ATC ACC ATC AAA GAA AGG CCG GAG ACA TTT TCA AGG ATA | 72 |
| 5wf-009 1NO | AAA ATT TTA GCC TTT ATT TCA ACG TTT TTC TAC TAA TAG TAG TAA AAA GGT GGC ATC AAT | 60 |
| 5wf-010 2AB | GTC TGA AAC ACG ACC AGT AAT AAA TGC GCG AAC TGA TAG CAC CAC CAG CAG AAG ATA AAA CAG A | 64 |
| 5wf-011 2CD | AGG GAT TTG TGT TTT TAT AAT CAG CGC AAA TTA ACC GTT GCT TGC CTG AGT AGA AGG CTC AAT C | 64 |
| 5wf-012 2EF | AAG GAA GGT GGC AAG TGT AGC GGT TTA ATG CGC CGC TAC ATA TAA CGT GCT TTC CTC CGA TTA A | 64 |
| 5wf-013 2GH | CAT TAA TTA AAC CTG TCG TGC CAG AGG CGA AAA TCC TGT TAT AAA TCA AAA GAA TAT GGC GAG A | 64 |
| 5wf-014 2IJ | CGT AAC CGA TCG GCC TCA GGA AGA GTT GGG TAA CGC CAG GCC AGT GCC AAG CTT GCC TAA CTC A | 64 |
| 5wf-015 2KL | GAG AGA TCT GGA GCA AAC AAG AGA CAA TCA TAT GTA CCC CAG ATT GTA TAA GCA AAG GGC GCA T | 64 |
| 5wf-016 2MN | GCG GGA GAT AGA ACC CTC ATA TAT AAG ATT CAA AAG GGT GTA TGA TAT TCA ACC GTC TAT TTT T | 64 |
| 5wf-017 2OP | CTA AAG TAG GAA GTT TCA TTC CAT TTT GGG GCG CGA GCT GGC ATT AAC ATC CAA TAA TAC TTT T | 64 |
| 5wf-018 3AB | GGT GAG GCG GCT ATT AGT CTT TAA AGG GAC ATT CTG GCC AAT ACC TAC ATT TTG ACA ACT CAA A | 64 |

| | | |
|---|---|---|
| 5wf-019 3C | CTA TCG GCA AGA GTC TGT CCA TCA TGA GGC CA | 32 |
| 5wf-020 3D | CCG AGT AAG GAG CTA AAC AGG AGG CGT TAG AA | 32 |
| 5wf-021 3E | TCA GAG CGA CCA CAC CCG CCG CGC CAC GCT GC | 32 |
| 5wf-022 3F | GCG TAA CCG GAA AGC CGG CGA ACG GCC CGA GA | 32 |
| 5wf-023 3G | TAG GGT TGG CTG GTT TGC CCC AGC CTG CAT TA | 32 |
| 5wf-024 3H | ATG AAT CGG TGC CTA ATG AGT GAG ATG CCT GC | 32 |
| 5wf-025 3I | AGG TCG ACG CTG CAA GGC GAT TAA TCG CAC TC | 32 |
| 5wf-026 3J | CAG CCA GCT CAC GTT GGT GTA GAT TAT TTA AA | 32 |
| 5wf-027 3K | TTG TAA ACC GTA AAA CTA GCA TGT ATC GAT GA | 32 |
| 5wf-028 3L | ACG GTA ATA TGC CGG AGA GGG TAG TCT AGC TG | 32 |
| 5wf-029 3M | ATA AAT TAG AGT AAT GTG TAG GTA TTT AAA TG | 32 |
| 5wf-030 3N | CAA TGC CTA CAT TAT GAC CCT GTA AAT CAT AC | 32 |
| 5wf-031 3OP | AGG CAA GGT TTA GCT ATA TTT TCA ATA ACA GTT GAT TCC CGC TCA ACA TGT TTT AAA TAT GCA A | 64 |
| 5wf-032 4A | TTT TGA ATG GTC AGT ATT AAC ACC GCC TGC AAC AGT GCC A | 40 |
| 5wf-033 4B | CTT GCT GGT AAT ATC CAA AAA CGC TCA TGG AAA CAG AGA TAG AAC CCT GAC AAT AT | 56 |
| 5wf-034 4G | CGG TCC ACA GTG TTG TTC CAG TTT ATT TAG AGC TTG ACG G | 40 |
| 5wf-035 4H | AGC CTG GGG CCA ACG CGC GGG GAG GCA GCA AG | 32 |
| 5wf-036 4I | GGG GAT GTT CTA GAG GAT CCC CGG AAG TGT AA | 32 |
| 5wf-037 4J | GTT AAT ATT TTG TTA AAC CGT AAT GGG ATA GGT TTC CGG CAC CGC TTC GGC GAA AG | 56 |
| 5wf-038 4O | ATA ACC TGC AAA GAA TTA GCA AAA ATC GGT TGT ACC AAA A | 40 |
| 5wf-039 4P | CTT AGA GCT TAA TTG CTG AAT ATA ATG CTG TAA ATT CTG CGA ACG AGT AAT GGT CA | 56 |
| 5wf-040 5-6A | AAG GAA TTG TCA GTT GGC AAA TCA AAG AAT ACG TGG CAC ATC TGA CCT GAA AGC GTA CAG TTG A | 64 |
| 5wf-041 5-6B | ATT ACC GCT AAT TTT AAA AGT TTG TTT GCC CGA ACG TTA TCA GCC ATT GCA ACA GGA GAA CAA T | 64 |
| 5wf-042 5F | TCG GAA CCC TAA AGG GAG CCC CCG GGA ACA AG | 32 |
| 5wf-043 5G | AGT CCA CTT GGC CCT GAG AGA GTT AGG CGG TT | 32 |
| 5wf-044 5H | TGC GTA TTA CGA GCC GGA AGC ATA GTA CCG AG | 32 |
| 5wf-045 5I | CTC GAA TTC GCT ATT ACG CCA GCT TGG TGC CG | 32 |
| 5wf-046 5J | GAA ACC AGA ACA AAC GGC GGA TTG AAT TCG CA | 32 |
| 5wf-047 5-6N | TAA AGC CTA ATC CCC CTC AAA TGC TCA TAA ATA TTC ATT GCA GAG CAT AAA GCT AAT TAA GCA A | 64 |
| 5wf-048 5-6O | TCA AAA AGT AGT CAG AAG CAA AGC TTA GAT ACA TTT CGC AAG ATT TAG TTT GAC CAG GAT TGC A | 64 |
| 5wf-049 6G | TCA CCG CCA TTA AAG AAC GTG GAC GCA CTA AA | 32 |
| 5wf-050 6H | CAC CGA CTC CAA AGA CAA AAG GGC GAT TAA GA | 32 |
| 5wf-051 6I | GGC CTC TTC GTA ATC ATG GTC ATA CAA TTC CA | 32 |
| 5wf-052 6J | TCC GTG GGG CAA AGC GCC ATT CGC TCG GTG CG | 32 |
| 5wf-053 6K | TTA AAT TTT TGT TAA ATC AGC TCA TCG GAT TC | 32 |
| 5wf-054 7A | CGC TGA GAG CCA GCA GCA AAT GAA AAA TCT AAC CTC AAT C | 40 |

| | | |
|---|---|---|
| 5wf-055 7B | AAT ATC TGG AGG AAG GTT ATC TAA AAC TCG TAT TAA ATC CAG TAA CAT TAT CAT TT | 56 |
| 5wf-056 7G | GTC GAG GTG CCG TAA ATC CAA CGT CAA AGG GCA GAC GGG C | 40 |
| 5wf-057 7H | AAC AGC TGT TTC TTT TCA CCA GTG AAT TGT TA | 32 |
| 5wf-058 7I | TCC GCT CAG CTG TTT CCT GTG TGA CTG TTG GG | 32 |
| 5wf-059 7J | AAG GGC GAC ATT CAG GCT GCG CAA AGC GAG TAA CAA CCC GTT TTT TAA CCA ATA GG | 56 |
| 5wf-060 7O | CAA TAC TGC GGA ATC GTT TAA ACA GTT CAG AAT TTA CCC T | 40 |
| 5wf-061 7P | GAC TAT TAA TTA AGA GGA AGC CCG TCC TTT TGA TAA GAG GTC ATT TTT GCG GAT GG | 56 |
| 5wf-062 8-9A | AGA AAA CTA CCT CAA ATA TCA AAC AGC ATC ACC TTG CTG ATT TTC AAA TAT ATT TTA GAA CGC G | 64 |
| 5wf-063 8B | AAA GAA ACT TAC AAA CAA TTC GAC AAT ATC TTT AGG AGC ACT AAC AAC | 48 |
| 5wf-064 8C | CAT ATT CCC ACC AGA AGG AGC GGA TGC GGA AC | 32 |
| 5wf-065 8D | CAA AAT TAT GGA AGG GTT AGA ACC ATT ATC AT | 32 |
| 5wf-066 8E | AAA TTG CGT TTG CAC GTA AAA CAG TAC CAT AT | 32 |
| 5wf-067 8F | GAA AAA CCG TCT ATC AAA TCA AGT TTT TTG GGA AAT AAA G | 40 |
| 5wf-068 8K | CTA ACG GAA ACG CCA TCA AAA ATA ATC AAC ATT AAA TGT G | 40 |
| 5wf-069 8L | TAC GAG GCA CAA CAT TAT TAC AGG AAA ACG AA | 32 |
| 5wf-070 8M | CCC TCG TTA TAG TAA GAG CAA CAC AAA GGA AT | 32 |
| 5wf-071 8N | ATA GCG TCT ACC AGA CGA CGA TAA TAT CAT AA | 32 |
| 5wf-072 8O | CAA ATA TCA ATC AAA AAT CAG GTC AAC GAG AAT GAC CAT ATA GAC TGG | 48 |
| 5wf-073 8-9P | ATT CGA GCA CAG GTC AGG ATT AGA GAG TAC CTT TAA TTG CAA AGA CTT | 48 |
| 5wf-074 9BC | TAA TAG ATT TAG AAG TAT TAG ACT GAA ACA GTA CAT AAA TAT TAC CTT | 48 |
| 5wf-075 9D | TTT TAA TGT GAT TAT CAG ATG ATG TTA TAC TTC TGA ATA AAG TTA CAA | 48 |
| 5wf-076 9E | AAT CGC GCA TTG CTT TGA ATA CCA TAG ATT TTC AGG TTT AAC GTC AGA | 48 |
| 5wf-077 9-10F | ATC ACC CAG GGC GAT GGC CCA CTA GAG AGA TAA CCC ACA ATT GAG CGC TAA TAT CAC GTG AAC C | 64 |
| 5wf-078 9-10J | CTA TTT CGG TAT AAA CAG TTA ATG CCT GTA GCC AGC TTT CAT TCG CGT CTG GCC TTC CCC CTG C | 64 |
| 5wf-079 9KL | ACG TTA ATT AGA AAG ATT CAT CAG CAG ATA CAT AAC GCC AAG AAC GAG | 48 |
| 5wf-080 9MN | TAG TAA ATC CTG ACG AGA AAC ACC AAA CCA AAA TAG CGA GGT AAT AGT | 48 |
| 5wf-081 9O | AAA ATG TTC AGA CGG TCA ATC ATA TAG CCG GAA CGA GGC GGC GTT TTA | 48 |
| 5wf-082 10-11A | CAA GAC AAA GTT AAT TTC ATC TTC TGA CCT AAA TTT AAT GTA AAT GCT | 48 |
| 5wf-083 10BC | GTG AGT GAT AAT ACA TTT GAG GAT TAG AGC CGT CAA TAG ATC CAA TCG | 48 |
| 5wf-084 10D | TGT TTG GAG CAA TTC ATC AAT ATA TAA CAA TTT CAT TTG ACA ATA TAT | 48 |

| | | |
|---|---|---|
| 5wf-085 10E | TGA ATA TAA TAA CGG ATT CGC CTG AGA GGC GAA TTA TTC AAT CCT GAT | 48 |
| 5wf-086 10KL | AAC TAA TGT TGA GAT TTA GGA ATA TCA GGA CGT TGG GAA GAA AAA TCT | 48 |
| 5wf-087 10MN | GCA AAA GAA GTG AAT AAG GCT TGC TGG GCT TGA GAT GGT TCC ACA TTC | 48 |
| 5wf-088 10O | ATG TTA CTA GGG AAC CGA ACT GAC AGT TTT GCC AGA GGG GAG GCT TTT | 48 |
| 5wf-089 10-11P | GAA CCA GAA CAC TCA TCT TTG ACC AAA AGA ATA CAC TAA ACC GGA AGC AAA CTC CAT TCA AAG C | 64 |
| 5wf-090 11B | GAT GCA AAA ATA GTG AAT TTA TCA GAC GCT GAG AAG AGT CAT AAC CTT | 48 |
| 5wf-091 11C | GCT TCT GTC AAA ATT AAT TAC ATT ACA AAC AT | 32 |
| 5wf-092 11D | CAA GAA AAA AAA GAA GAT GAT GAA TTT CAA TT | 32 |
| 5wf-093 11E | ACC TGA GCT ACA TCG GGA GAA ACA CAG TAA CA | 32 |
| 5wf-094 11F | GTA CCT TTC AAA GTC AGA GGG TAA GAA TTG AGT TAA GCC C | 40 |
| 5wf-095 11K | TTG AGT AAC AGT GCC CGA ACC TAT TAT TCT GAT GGC TCA T | 40 |
| 5wf-096 11L | TAT ACC AGA TGC GAT TTT AAG AAC CAT TGT GA | 32 |
| 5wf-097 11M | ATT ACC TTT AAT TTC AAC TTT AAT ACA AAG CT | 32 |
| 5wf-098 11N | GCT CAT TCT ACC CAA ATC AAC GTA AGA ACC GG | 32 |
| 5wf-099 11O | ATA TTC ATC AAC TTT GAA AGA GGA GTG TCG AAA TCC GCG ACC TGC TCC | 48 |
| 5wf-100 12A | ACT ATA TGG TTT GAA ATA CCG ACC GTG TGA TAA ATA AGG C | 40 |
| 5wf-101 12B | AAA TCG TCG CTA TTA ACG ATA GCT TAG ATT AAA AAT CAT AGG TCT GAG GTT ATA TA | 56 |
| 5wf-102 12G | AGA AAC GCA ATA ATA AGA GCA AGA AAC TGA ACA CCC TGA A | 40 |
| 5wf-103 12H | AAG GCC GGA AAG ACA CCA CGG AAT CAT ATA AA | 32 |
| 5wf-104 12I | CAG AGC CAA AAC GTC ACC AAT GAA CCA TTA GC | 32 |
| 5wf-105 12J | AAC ATG AAA GTA TTA ATA ACG GGG TCA GTG CCC CAC CCT CAG AGC CGC GCC ACC CT | 56 |
| 5wf-106 12O | GAT AAA TTC AGA TGA ACG GTG TAC AAG AGT AAT CTT GAC A | 40 |
| 5wf-107 12P | CTA CGA AGG CAC CAA CCT AAA ACG AAA GAG GCC CCA GCG ATT ATA CCA CAT CGC CT | 56 |
| 5wf-108 13-14A | GAG CCA GTT GTA ATT TAG GCA GAG CTC CGG CTT AGG TTG GAG ACT ACC TTT TTA ACG CAT TTT C | 64 |
| 5wf-109 13-14B | CCC TTA GAT TTA CGA GCA TGT AGA TAA TAT CCC ATC CTA AAT CCT TGA AAA CAT AGT TAA TTT T | 64 |
| 5wf-110 13F | GGA AGC GCA TTA GAC GGG AGA ATT AAC AAT GA | 32 |
| 5wf-111 13G | AAT AGC AAT ACA TAA AGG TGG CAA AAG TTT AT | 32 |
| 5wf-112 13H | TTT GTC ACA CCA GTA GCA CCA TTA ACC ATC GA | 32 |
| 5wf-113 13I | TAG CAG CAC GCC ACC CTC AGA ACC CAC CAG AA | 32 |
| 5wf-114 13J | CCA CCA CCT ACT GGT AAT AAG TTT GAG GCT GA | 32 |
| 5wf-115 13-14N | AAA AGG AGG AAA ATC TCC AAA AAA CTG GCT GAC CTT CAT CAG ACC AGG CGC ATA GGA AGG CTC C | 64 |
| 5wf-116 13-14O | ACA AAG TAG CCG CTT TTG CGG GAT TTG CAG GGA GTT AAA GCA ACG GAG ATT TGT ATA GCG CGA A | 64 |

| | | |
|---|---|---|
| 5wf-117 14G | AAA ATA CAT AGC TAT CTT ACC GAA AAA AAC AG | 32 |
| 5wf-118 14H | GCA AAA TCA ATC AAT AGA AAA TTC AAA CGT AG | 32 |
| 5wf-119 14I | CTC AGA GCC CGT AAT CAG TAG CGA TAG AGC CA | 32 |
| 5wf-120 14J | CAG GAG TGA GAG CCG CCG CCA GCA CCG CCT CC | 32 |
| 5wf-121 14K | GAC TCC TCA AGA GAA GGA TTA GGA TGA TGA TA | 32 |
| 5wf-122 15A | GTT AAA TAA GAA TAA ACA CCG GAA TCA TAA TTT TTA ACA A | 40 |
| 5wf-123 15B | CGC CAA CAA ATA AGA GAA TAT AAA TCC TGA ACA AGA AAA AAA CCA ATC AAT AAT CG | 56 |
| 5wf-124 15G | ACA GAG AGA ATA ACA TGC CCT TTT TAA GAA AAT ACG CAG T | 40 |
| 5wf-125 15H | ATG TTA GCA TAT GGT TTA CCA GCG TGA GCC AT | 32 |
| 5wf-126 15I | TTG GGA ATC AGA ATC AAG TTT GCC AGA GCC AC | 32 |
| 5wf-127 15J | CAC CGG AAT TGA CAG GAG GTT GAG CGT CAT ACA TGG CTT TTT AGC GGG GTT TTG CT | 56 |
| 5wf-128 15O | ATA ATT TTT TCA CGT TCC TTT AAT TGT ATC GGA TTC GGT C | 40 |
| 5wf-129 15P | GCT GAG GCC GTC ACC CTC AGC AGC TTT CCA TTA AAC GGG TAA AAT ACG TAA TGC CA | 56 |
| 5wf-130 16AB | TAG ATA AGG TAC CGA CAA AAG GTA ATT GAG AAT CGC CAT AAC TAG AAA AAG CCT GTT TAG TAT C | 64 |
| 5wf-131 16C | GGA ATC ATG CTG TCT TTC CTT ATC ATC AAC AA | 32 |
| 5wf-132 16D | CGG GAG GTT ACC GCG CCC AAT AGC TCA TCG TA | 32 |
| 5wf-133 16E | AGA GCC TAT TTG AAG CCT TAA ATC CCG ACT TG | 32 |
| 5wf-134 16F | CAG CCT TTA TTT GCC AGT TAC AAA GTC TTT CC | 32 |
| 5wf-135 16G | CTC CTT ATG TAA GCA GAT AGC CGA GAA AAT AG | 32 |
| 5wf-136 16H | CAC AAC ATG GGC GCC AGG GTG GTT ATT GCC CT | 32 |
| 5wf-137 16I | CCG GAA CCT TTA GCG TCA GAC TGT ATC ACC GT | 32 |
| 5wf-138 16J | CCA GTA AGG CAG GTC AGA CGA TTG CAA AAT CA | 32 |
| 5wf-139 16K | GCC ACC CTC AGT ACC AGG CGG ATA TTA CCG TT | 32 |
| 5wf-140 16L | ACA CTG AGC AGA ACC GCC ACC CTC TTA GTA CC | 32 |
| 5wf-141 16M | CCA GAC GTT TTC GTC ACC AGT ACA GTA CCG TA | 32 |
| 5wf-142 16N | TGC GAA TAT AGT AAA TGA ATT TTC TCG TCT TT | 32 |
| 5wf-143 16OP | TGA GGA AGG AAA GAC AGC ATC GGA CAC GCA TAA CCG ATA TTT TAT CAG CTT GCT TTA AAG GAA T | 64 |
| 5wf-144 17AB | ATA TGC GTT CAA CAG TAG GGC TTA AAG TAA TTC TGT CCA GCA GAA CGC GCC TGT TTA TTC CAA G | 64 |
| 5wf-145 17CD | AAC GGG TAC AAG CCG TTT TTA TTT AAG CAA ATC AGA TAT ACG TTT TAG CGA ACC TCA AGA TTA G | 64 |
| 5wf-146 17EF | TTG CTA TTA CCA ACG CTA ACG AGC ATA AAC AGC CAT ATT AGT TTA ACG TCA AAA ATA CAA AGT T | 64 |
| 5wf-147 17GH | ACC AGA AGC CAA AAG AAC TGG CAT GAC ATT CAA CCG ATT GTC ATT AAA GGT GAA TTA GCG CGT T | 64 |
| 5wf-148 17IJ | TTC ATC GGG CCA TCT TTT CAT AAT GCC TTG ATA TTC ACA AAG CGC AGT CTC TGA ATA GTG CCG T | 64 |
| 5wf-149 17KL | CGA GAG GGA CCG TAC TCA GGA GGT AGA ACC GCC ACC CTC ACC CAA TAG GAA CCC ATA ACT ACA A | 64 |
| 5wf-150 17MN | CGC CTG TAA CGA TCT AAA GTT TTG TGT ATG GGA TTT TGC TAT AGA AAG GAA CAA CTC GAG GTG A | 64 |

| | | |
|---|---|---|
| 5wf-151 17OP | ATT TCT TAG ACA ACA ACC ATC GCC ACG AGG GTA GCA ACG GCT TTG AGG ACT AAA GAC TTT TTC A | 64 |
| 5wf-152 18A | AAT AAA CAT TTT AGT ATA AAG CCA ACG CTA TAC AAA TTC TTA CC | 44 |
| 5wf-153 18BC | ATC CGG TAT TTT ACT CAT CGA GAA CAA GTT AAA CCA AGT ACC GCT TTT ACA TGT TCA GCT AAT GAC GAC GAC | 72 |
| 5wf-154 18DE | AAT CCA AAT TTT TTT TAT CCT GAA TCT TTT GCA CCC AGC TAC AAT TTT TTC TAA GAA CGC GAG GGA AGG CTT | 72 |
| 5wf-155 18FG | AAG GTA AAT TTT AAT AAT AAC GGA ATA CGA AAC CGA GGA AAC GCT TTT TAA GAA ACG ATT TTT TTT TAT CCC | 72 |
| 5wf-156 18HI | ATC CTC ATT TTT CCC CTT ATT AGC GTT TCA TTT TCG GTC ATA GCT TTT TAT TGA CGG AAA TTA TAG GGA GGG | 72 |
| 5wf-157 18JK | ACC CTC ATT TTT CCG GAA TAG GTG TAT CTT GAT ATA AGT ATA GCT TTT TAA AGC CAG AAT GGA AAC AAA TAA | 72 |
| 5wf-158 18LM | TTC AAC AGT TTT CCT CAT AGT TAG CGT AGC ATT CCA CAG ACA GCT TTT TTT CAG GGA TAG CAA GGA GCC ACC | 72 |
| 5wf-159 18NO | TAG TTG CGC CGA CAA TAA CAG CTT GAT ACC GAT TTT TTT CAG CGG AGT GAG AAA ACA ACT | 60 |

**Table S-1.** Sequences of the 159 complementary oligos.

The DNA nanostructure was made by mixing together 2 μL M13mp18 DNA (10 nM), 5μL oligonucleotide mix (each oligo 200 nM), 2 μL origami buffer 10x (20 mM MgCl2, 100 mM Tris-HCl, pH=7.6), and 11 μL deionised MilliQ water. EDTA was omitted from the origami buffer for the whole of this study (normally included at 1mM), so as not to interfere with the bacterial populations. The mixture was annealed from 85 to 25 °C at a rate of −1.0 °C/min. After annealing, excess oligonucleotides were removed using a Micro Biospin column (Bio-Rad) packed with Sephacryl S-300 (GE Healthcare).

##### 1.2 Modifications to the 5-well frames

The following oligonucleotides were modified to carry aptamers that can bind *E.coli* and *B.subtilis*[2]. The aptamers sequence is CAT ATC CGC GTC GCT GCG CTC AGA CCC ACC ACC ACG CAC C (in red in the table below).

| Sequence Name | Sequence | Bases |
|---|---|---|
| 5wf-009 1NO-apt | AAA ATT TTA GCC TTT ATT TCA ACG TTT TTC TAC TAA TAG TAG TAA AAA GGT GGC ATC AAT TTT TT C ATA TCC GCG TCG CTG CGC TCA GAC CCA CCA CCA CGC ACC | 105 |
| 5wf-015 2KL-apt | GAG AGA TCT GGA GCA AAC AAG AGA CAA TCA TAT GTA CCC CAG ATT GTA TAA GCA AAG GGC GCA TTT TTT CAT ATC CGC GTC GCT GCG CTC AGA CCC ACC ACC ACG CAC C | 109 |

| | | |
|---|---|---|
| 5wf-159 18NO-apt | TAG TTG CGC CGA CAA TAA CAG CTT GAT ACC GAT TTT TTT CAG CGG AGT GAG AAA ACA ACT TTT TT C ATA TCC GCG TCG CTG CGC TCA GAC CCA CCA CCA CGC ACC | 105 |
| 5wf-001 1A-apt | TAA AAA TAC CGA ACG ACC TAA AAC ATC GCC ATG CAG ATT CTT TTT CAT ATC CGC GTC GCT GCG CTC AGA CCC ACC ACC ACG CAC C | 85 |
| 5wf-023 3G-apt | TAG GGT TGG CTG GTT TGC CCC AGC CTG CAT TAT TTT T CA TAT CCG CGT CGC TGC GCT CAG ACC CAC CAC CAC GCA CC | 77 |
| 5wf-043 5G - apt | AGT CCA CTT GGC CCT GAG AGA GTT AGG CGG TTT TTT T CA TAT CCG CGT CGC TGC GCT CAG ACC CAC CAC CAC GCA CC | 77 |
| 5wf-045 5I - apt | CTC GAA TTC GCT ATT ACG CCA GCT TGG TGC CGT TTT T CA TAT CCG CGT CGC TGC GCT CAG ACC CAC CAC CAC GCA CC | 77 |
| 5wf-075 9D-apt | TTT TAA TGT GAT TAT CAG ATG ATG TTA TAC TTC TGA ATA AAG TTA CAA TTT TT C ATA TCC GCG TCG CTG CGC TCA GAC CCA CCA CCA CGC ACC | 93 |
| 5wf-111 13G-apt | AAT AGC AAT ACA TAA AGG TGG CAA AAG TTT ATT TTT T CA TAT CCG CGT CGC TGC GCT CAG ACC CAC CAC CAC GCA CC | 77 |
| 5wf-113 13I - apt | TAG CAG CAC GCC ACC CTC AGA ACC CAC CAG AAT TTT T CA TAT CCG CGT CGC TGC GCT CAG ACC CAC CAC CAC GCA CC | 77 |
| 5wf-136 16H-apt | CAC AAC ATG GGC GCC AGG GTG GTT ATT GCC CTT TTT T CA TAT CCG CGT CGC TGC GCT CAG ACC CAC CAC CAC GCA CC | 77 |
| 5wf-138 16J - apt | CCA GTA AGG CAG GTC AGA CGA TTG CAA AAT CAT TTT T CA TAT CCG CGT CGC TGC GCT CAG ACC CAC CAC CAC GCA CC | 77 |
| 5wf-152 18A-apt | AAT AAA CAT TTT AGT ATA AAG CCA ACG CTA TAC AAA TTC TTA CCT TTT T CA TAT CCG CGT CGC TGC GCT CAG ACC CAC CAC CAC GCA CC | 89 |
| 5wf-079 9KL-apt | ACG TTA ATT AGA AAG ATT CAT CAG CAG ATA CAT AAC GCC AAG AAC GAG TTT TT C ATA TCC GCG TCG CTG CGC TCA GAC CCA CCA CCA CGC ACC | 93 |

**Table S-2.** Sequences of the 14 aptamer-modified oligos.

The following oligonucleotides were functionalised with Alexa 647 molecules.

| | | |
|---|---|---|
| 5wf-025 3I 647 | /5Alex647N/TTT TTA GGT CGA CGC TGC AAG GCG ATT AAT CGC ACT C | 37 |
| 5wf-126 15I 647 | /5Alex647N/TTT TTT TGG GAA TCA GAA TCA AGT TTG CCA GAG CCA C | 37 |
| 5wf-083 10BC 647 | /5Alex647N/TTT TTG TGA GTG ATA ATA CAT TTG AGG ATT AGA GCC GTC AAT AGA TCC AAT CG | 53 |
| 5wf-081 9O 647 | /5Alex647N/TTT TTA AAA TGT TCA GAC GGT CAA TCA TAT AGC CGG AAC GAG GCG GCG TTT TA | 53 |

**Table S-3.** Sequences of the 4 Alexa 647-modified oligos.

The following oligonucleotides were functionalised to carry biotin:

| | | |
|---|---|---|
| 5wf-057 7H-bio | AAC AGC TGT TTC TTT TCA CCA GTG TTT TT/3Bio/ | 29 |
| 5wf-066 8E-bio | AAA TTG CGT TTG CAC GTA AAA CAG TTT TT/3Bio/ | 29 |
| 5wf-096 11L-bio | TAT ACC AGA TGC GAT TTT AAG AAC TTT TT/3Bio/ | 29 |

| | | |
|---|---|---|
| 5wf-020 3D-bio | /5Biosg/TTT TTG GAG CTA AAC AGG AGG CGT TAG AA | 29 |
| 5wf-028 3L-bio | /5Biosg/TTT TTA TGC CGG AGA GGG TAG TCT AGC TG | 29 |
| 5wf-070 8M-bio | /5Biosg/TTT TTA TAG TAA GAG CAA CAC AAA GGA AT | 29 |
| 5wf-092 11D-bio | /5Biosg/TTT TTA AAA GAA GAT GAT GAA TTT CAA TT | 29 |
| 5wf-104 12I-bio | /5Biosg/TTT TTA AAC GTC ACC AAT GAA CCA TTA GC | 29 |
| 5wf-133 16E-bio | /5Biosg/TTT TTT TTG AAG CCT TAA ATC CCG ACT TG | 29 |
| 5wf-141 16M-bio | /5Biosg/TTT TTT TTC GTC ACC AGT ACA GTA CCG TA | 29 |

**Table S-4.** Sequences of the 10 biotin-modified oligos.

#### Section 2. Imaging of the nanostructures

##### 2.1 Atomic Force Microscopy (AFM)

Origami tiles were diluted ten times in origami buffer and 25 µl of the sample were deposited on freshly cleaved mica and incubated at room temperature for 10 minutes. The samples were then washed 5 times with 1 ml of origami buffer. The nanostructures were imaged in origami buffer, using a Dimension FastScan AFM microscope (Bruker). The probes used were FastScan-D probes (Bruker), with a resonant frequency of 90 kHz, a spring constant of 0.21 $Nm^{-1}$ and a nominal tip radius of 8 nm.

DNA origami tiles were incubated with 0.17 µM Streptavidin (Sigma Aldrich) for 5 minutes at room temperature, after which they were filtered using a Millipore filter (Millipore, MA, USA) unit with molecular weight cut-off (MWCO) of 100 KDa to remove free protein. The tiles were then prepared for AFM imaging as described above.

To load the streptavidin functionalised origami tiles with the antimicrobial enzyme, the tiles were incubated with 1mg/ml biotinylated lysozyme (Chicken Lysozyme protein, Egg whites, GeneTex) for 10 minutes at room temperature after which they were filtered and imaged as above.

##### 2.2 AFM data analysis

Images of all datasets were plane-fitted using the speed-optimised plane correction function of the SPIP software (Image Metrology A/S, Hørsholm, Denmark), which fits each line in the horizontal axis to a polynomial equation. SPIP was also used for calculation of the volumes of proteins attached to the DNA origami tiles. The “inspection window” feature of SPIP was used to zoom into individual tiles and then the “circular area of interest” tool was used to allow the software to calculate only the volume of the protein rather than that of the whole tile, according to the following equation:

$$Z_{\text{net volume}} = Z_{\text{material volume}} - Z_{\text{void volume}}$$

where $Z_{\text{material volume}}$ is the volume of all pixels inside the shape’s contour with a Z value greater than zero:

$$Z_{\text{material volume}} = \sum_{\{Z(x,y)\in \text{shape} \mid Z \geq 0\}} Z(x,y)\,dxdy$$

where dx and dy are the point spacings in the X and Y directions of the image, respectively. $Z_{\text{void volume}}$ is the volume of all pixels inside the shape’s contour with a Z value lower than or equal to zero:

$$Z_{\text{void volume}} = \sum_{\{Z(x,y)\in \text{shape} \mid Z \leq 0\}} Z(x,y)\,dxdy$$

where dx and dy are the point spacing in the X and Y directions of the image, respectively.

For cross-sections of sample features, tile dimensions measurements, as well as for the 3D rendering of the images, Nanoscope 1.9 software (Bruker) was used. Volume histograms were drawn with bin widths chosen according to Scott’s equation, using GraphPad Prism.

“Theoretical” molecular volumes of proteins based on molecular mass were calculated using the equation by Schneider et al[3]:

$$V=(M_0/N_0)(V_1 + dV_2)$$

where $M_0$ is the molecular mass, $N_0$ is Avogadro’s number, $V_1$ and $V_2$ are the partial specific volumes of protein and water, respectively, and d is the extent of protein hydration. The partial specific volume of a typical protein ($V_1$) is considered to be 0.74 $cm^3g^{-1}$, and the extent of protein hydration (d) has been estimated to be 0.4 g of water/g of protein. The partial specific volume of water ($V_2$) is 1 $cm^3g^{-1}$

##### 2.3 direct Stochastic Optical Reconstruction Microscopy (dSTORM)

**Fluorescence microscopy experiments**

The fluorescence microscopy experiments performed on the origami structures were conducted on a custom-built microscope based on an Olympus (Center Valley, PA) IX-73 frame with a 100x 1.49 NA oil objective lens (Olympus UAPON100XOTIRF) and a 647-nm laser (MPB Communications Inc. VFL-P-300-647-OEM1-B1). The samples were imaged in total internal reflection fluorescence (TIRF) mode and images were relayed onto the camera (Andor iXon Ultra 897) by a 1.3x magnification Cairn Twincam image splitter (the second port of the image splitter was not used during these experiments).

**dSTORM on origami structures**

16000 frames at a 20 ms exposure time with an EM gain of 200 over a 256x256 pixel region were acquired for the dSTORM reconstructions. The pixel size in the image plane was measured to be 118 nm. The raw single molecule data sets were reconstructed using ThunderSTORM[4], and visualized as averaged shifted histograms with a magnification factor of 10.

The peak-to-peak distance between the fluorophores tethered to the origami structures was measured by taking a cross-sectional profile in Fiji/ImageJ[5] between two bright spots in different regions of interest in the reconstructed image, and using a custom MATLAB (Natick,MA) script to measure the average distance between two peaks.

#### Section 3. Imaging of bacterial populations

##### 3.1 Sample preparation for SIM

Each one of the bacterial strains was grown overnight ($OD_{600}$ of ~1 in the case of *E. coli* and ~0.6 in the case of *B. subtilis*). Prior to SIM imaging the cultures were spun down and washed three times in origami buffer. The bacteria were then resuspended in 100 µl origami buffer. In the case of *B. subtilis*, the bacterial pellet was resuspended in 100 µl origami buffer containing 1 µg/ml Nile red dye (Sigma-Aldrich, 72485). This step was not required for *E. coli,* as the BL21(DE3) *E.coli* cells used in this experiment had been transformed with pUC19GFP plasmid, and strongly express GFP.

Subsequently, 10 µl of bacterial suspension were mixed with 10µl of DNA origami (final concentration ~10 nM) and incubated with shaking at room temperature for 15 mins. The bacteria were gently centrifuged and resuspended in origami buffer to remove excess or unbound origami tiles. 2µl of the sample were deposited on a glass coverslip and an agarose pad was positioned over the sample to prevent the bacteria from moving. Another coverslip was positioned on top to minimise drying of the agarose pads.

**3.2 Structured Illumination Microscopy (SIM)**

Images of the sample were collected using 3-color SIM for optical sectioning[6]. A ×60/1.2 NA water immersion lens (UPLSAPO 60XW, Olympus) focused the structured illumination pattern onto the sample, and the same lens was also used to capture the fluorescence emission light before imaging onto an sCMOS camera (C11440, Hamamatsu). The wavelengths used for excitation were 488 nm (iBEAM-SMART-488, Toptica), 561 nm (OBIS 561, Coherent), and 640 nm (MLD 640, Cobolt). Images were acquired using custom SIM software described previously[7].

Large fields of view:

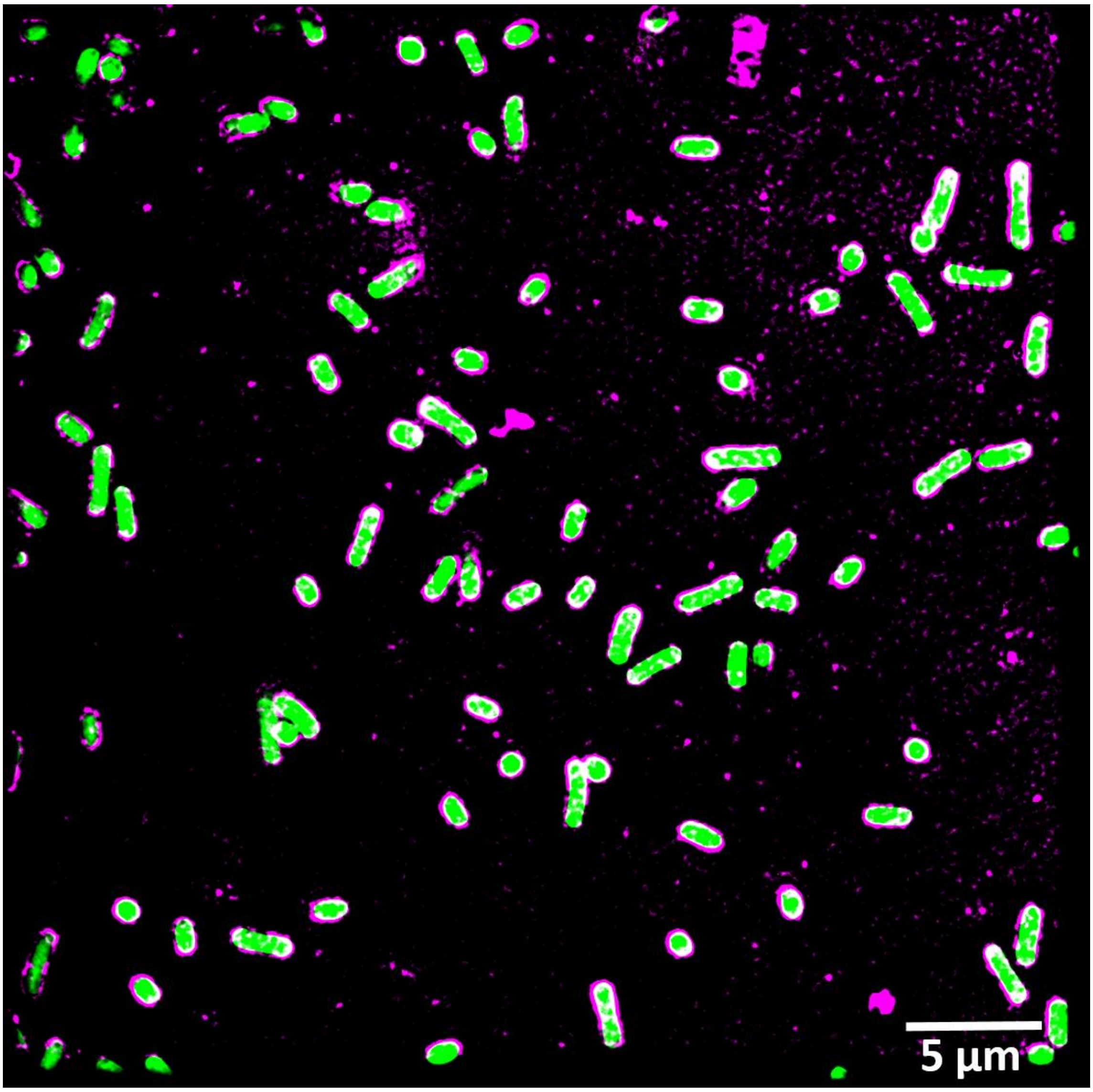


**Figure S.1.** Large field of view (42x42 μm) of E. coli (in green) decorated with DNA origami (in magenta). Overlap of the two colours is shown in white.

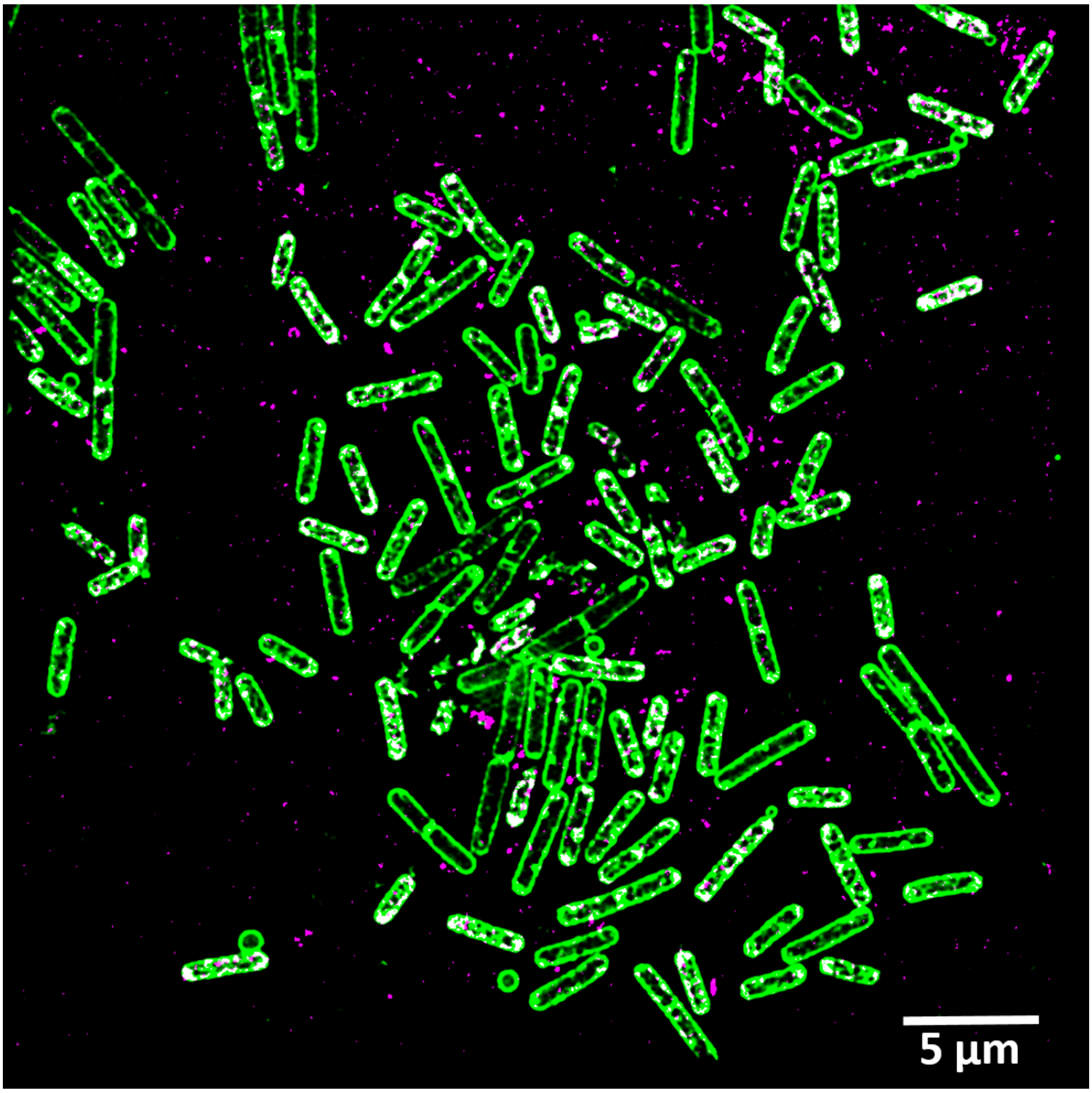


**Figure S.2** Large field of view (42x42 µm) of B. subtilis (in green) decorated with DNA origami (in magenta). Overlap of the two colours is shown in white.

Automated analysis of SIM images was performed with a custom-made MatLab routine. 2-colour SIM images contains information about bacterium body (green colour) and DNA-origami (magenta colour). The code defines coverage percentage of DNA origami as ratio between number of "overlapped" pixels (pixels where both colors have non-zero values of intensity) and number of all pixels corresponding to single bacterium (green color).

SIM imaging experiments were repeated three times, each time five fields of view were analysed for overlap for each strain, with ~ 825 single bacteria for *E. coli* and ~750 single bacteria for *B. subtilis* analysed in total.

#### Section 4. Bacterial growth curves

All bacterial cell culture studies were conducted using *E. coli* BL21(DE3), expressing GFP and *B. subtilis*. All experiments were conducted in LB medium, supplemented with carbenicillin (100 µg/ml) for *E. coli* and chloramphenicol (25 µg/ml) for *B. subtilis*. Bacterial starter cultures were grown overnight, and the bacteria were then diluted 1:100 into 150µl LB, and grown over 16 hours in a shaking plate reader, at 37°C, with measurements taken every 5 minutes, in the following conditions:

| ***E. coli*** | |
|---|---|
| **Sample 1** | **LB** |
| **Sample 2** | **LB + 10 nM DNA origami** |
| **Sample 3** | **LB + 0.3 µM free lysozyme** |
| **Sample 4** | **LB + 10 nM DNA origami carrying ~0.3µM lysozyme** |
| **Control 1** | **LB + 10 nM DNA origami w/o aptamers** |
| **Control 2** | **LB + Origami Buffer (10mM Tris, 2mM $MgCl_2$)** |

**Table S-5.** Experimental conditions for *E. coli*

| ***B. subtilis*** | |
|---|---|
| **Sample 1** | **LB** |
| **Sample 2** | **LB + 10 nM DNA origami** |
| **Sample 3** | **LB + 0.3 µM free lysozyme** |
| **Sample 4** | **LB + 10 nM DNA origami carrying ~0.3µM lysozyme** |
| **Control 1** | **LB + 10 nM DNA origami w/o aptamers** |
| **Control 2** | **LB + Origami Buffer (10mM Tris, 2mM $MgCl_2$)** |

**Table S-6.** Experimental conditions for *B. subtilis*

The OD values at 600nm were collected and used for the creation of growth curves. For each condition, 9 individual growth curves were analysed and averaged. Individual growth curves were fitted in MATLAB using the curve fitting toolbox, to a re-parameterised Gompertz growth model, to extract growth rates.

DNA origami carrying lysozyme were prepared as described in Section 2.1 and added to the samples where appropriate.

The growth rates for *E. coli* and *B. subtilis* grown in the presence of DNA origami without aptamers are presented below:

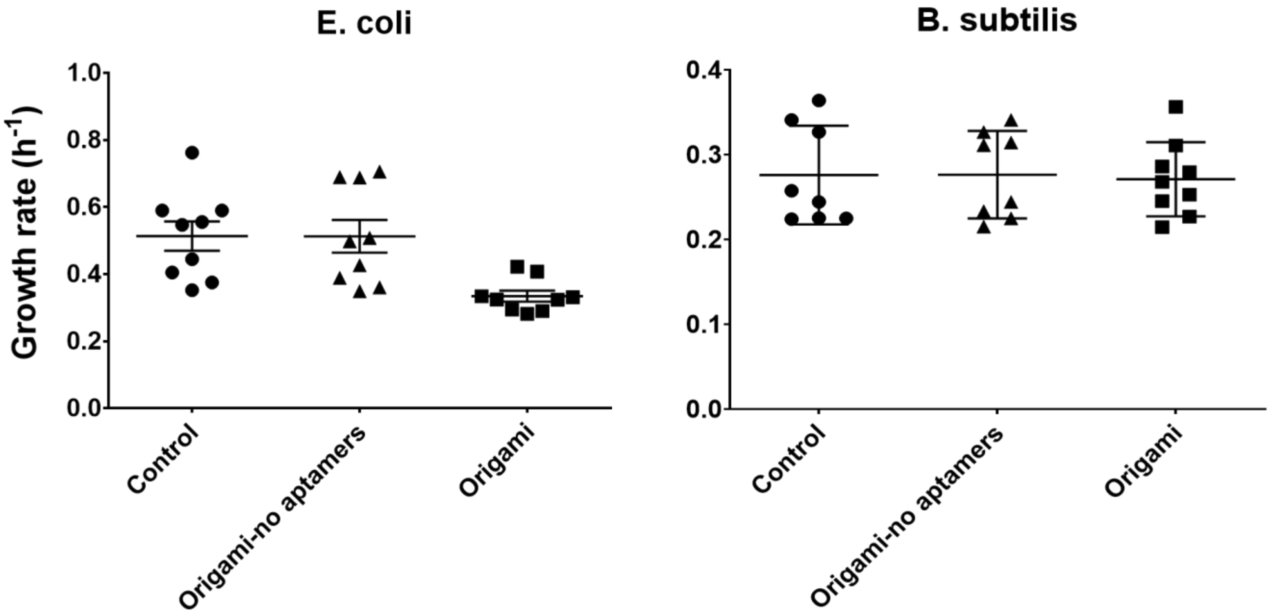


**Figure S.3:** Growth rates of *E. coli* and *B. subtilis* in the presence of DNA origami with and without aptamers.

#### Section 5. Cell viability assay

We used the CellTiter 96® AQueous One Solution Cell Proliferation Assay (Promega), to assess the effects of the DNA origami on mammalian cells. COS-7 cells were plated in a 96-well plate at concentration of 10,000 cells/well in 100µl of media (DMEM+10%FBS). 20µl of CellTiter 96® AQueous One Solution Reagent were added per well and the cells were incubated at 37°C for 2 hours in a humidified, 5% CO2 atmosphere. After 2 h, the absorbance at 490nm was measured, using a 96-well plate reader. The measurements were performed in triplicates.

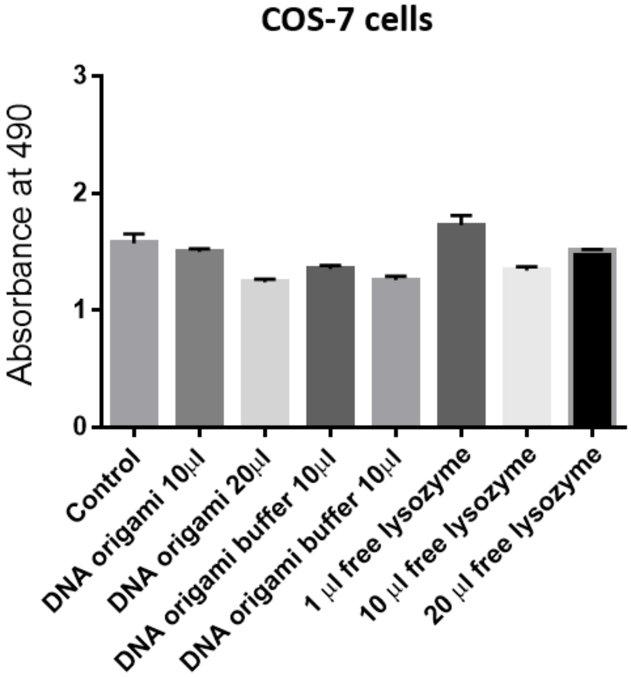


**Figure S.4:** Mammalian COS-7 cells are not affected by DNA origami
